# LRP1 Triggers Pro-inflammatory Cell-signaling in Response to Extracellular Tau Independently of the NMDA Receptor

**DOI:** 10.1101/2023.10.06.561299

**Authors:** Pardis Azmoon, Elisabetta Mantuano, Binita Poudel, Carlotta Zampieri, Steven L. Gonias

**Author notes:** Corresponding author: Steven L. Gonias. These authors contributed equally to this work (Pardis Azmoon, Elisabetta Mantuano).

## Abstract

In Alzheimer’s Disease (AD) and other neurodegenerative diseases, microtubule-associated protein Tau forms abnormal intracellular aggregates. The mechanisms by which Tau may promote AD progression remain incompletely understood. Injured and dying neurons release Tau into the extracellular spaces in the CNS. The released Tau may be taken up by receptors in the LDL Receptor gene family, including Low Density Lipoprotein Receptor-related Protein-1 (LRP1), which is expressed by microglia, astrocytes, and neurons. This process may be important for clearing Tau from extracellular spaces but may also promote the seeding of Tau aggregates in new cells. Our laboratory has shown that endocytosis of LRP1 ligands is coupled to the activation of cell-signaling and regulation of inflammation. Notably, different LRP1 ligands can induce either pro-inflammatory or anti-inflammatory responses, depending on the co-receptors that function with LRP1. Here, we demonstrate that in cultured macrophages, microglia, and astrocytes, extracellular Tau induces an LRP1-dependent pro-inflammatory response, characterized by NFκB activation and expression of pro-inflammatory cytokines. Unlike other LRP1 ligands that elicit anti-inflammatory responses, the response to Tau occurs independently of the NMDA receptor. When LRP1 is deleted or silenced, macrophages, microglia, and astrocytes do not respond to Tau, whereas when *Grin1* is deleted or the NMDA-R is pharmacologically inhibited, the responses remain unchanged. Because we have evidence that LRP1 in microglia expresses anti-inflammatory activity in response to ligands other than Tau, understanding the role of LRP1 in microglia and astrocytes *in vivo* in Alzheimer’s Disease and other neuroinflammatory processes is an important future goal.

**SIGNIFICANCE STATEMENT:** In Alzheimer’s Disease and other neurodegenerative diseases, microtubule-associated protein Tau forms abnormal intracellular aggregates that contribute to disease progression. When released extracellularly, Tau binds to the transmembrane receptor LRP1, expressed by diverse cells in the CNS. LRP1 has the unique ability to couple ligand uptake with activation of cell-signaling. We demonstrated that Tau binding to LRP1 activates pro-inflammatory responses in macrophages, microglia, and astrocytes, characterized by NFκB activation and cytokine release. This signaling occurs independently of the NMDA receptor, which distinguishes Tau from other LRP1 ligands. These results define a novel pathway by which extracellular Tau regulates neuroinflammation in Alzheimer’s Disease, providing new therapeutic opportunities that target LRP1 without interfering with NMDA-R functions.

## INTRODUCTION

Microtubule-associated protein Tau is an intracellular protein that forms abnormal oligomers and fibrils in Alzheimer’s Disease (AD) and in neurodegenerative diseases called Tauopathies (Lee et al., 2001; Congdon and Sigurdsson, 2018; Gao et al., 2018). Tau aggregation is regulated by gene polymorphisms, alternative splicing, phosphorylation, and other post-translational modifications. While the formation of neurofibrillary tangles through Tau aggregation is a key pathological hallmark, emerging evidence suggests that Tau’s contribution to AD extends beyond its aggregation state (Congdon and Sigurdsson, 2018; Gao et al., 2018).

Tau is released by neurons and other cells in the CNS, in extracellular vesicles (EVs) and independently of EVs, allowing for transfer of Tau between cells (Saman et al., 2012; Pooler et al., 2013; Asai et al., 2015; Evans et al., 2018; Merezhko et al., 2018; Brunello et al., 2020). Because Tau transfer between cells in the CNS may play a role in AD progression, identification of Low Density Lipoprotein Receptor-related Protein-1 (LRP1) as an endocytic receptor for extracellular Tau is highly significant (Rauch et al., 2020; Cooper et al., 2021). SORL1, which is structurally related to LRP1, also may mediate Tau uptake (Cooper et al., 2024).

LRP1 is a 600-kDa, type 1 transmembrane receptor for over 100 structurally diverse ligands, including apolipoprotein E, α_2_-macroglobulin (α_2_M), shed non-pathogenic cellular prion protein (S-PrP^C^), and proteins released from injured and dying cells (Strickland et al., 2002; Fernandez-Castaneda et al., 2013; Gonias and Campana, 2014; Mantuano et al., 2020). LRP1 ligands are internalized in clathrin-coated pits and usually transferred to lysosomes for degradation. A subset of LRP1 ligands trigger cell-signaling via mechanisms that require co-receptors, such as the NMDA Receptor (NMDA-R), Trk, p75^NTR^, and/or PrP^C^ (Bacskai et al., 2000; May et al., 2004; Martin et al., 2008; Shi et al., 2009; Mantuano et al., 2013; Stiles et al., 2013; Gonias and Campana, 2014; Krogh et al., 2014; Mattei et al., 2020). Assembly of different LRP1 co-receptors in response to different ligands may explain why the cell-signaling responses elicited by various LRP1 ligands are non-uniform (Gonias and Campana, 2014). LRP1 is expressed by neurons, astrocytes, oligodendrocytes, and microglia in the CNS and by macrophages throughout the body (Gonias and Campana, 2014; Auderset et al., 2016).

Binding of Tau to LRP1 is mediated by the Tau microtubule-binding domain (Rauch et al., 2020; Cooper et al., 2021). The Tau alternative splicing variant with four microtubule-binding repeats (2N4R) binds to LRP1 with higher affinity than the variant with three. Tau phosphorylation events, which exacerbate Tau aggregation (Alonso et al., 2001; Augustinack et al., 2002), apparently decrease LRP1-binding affinity (Cooper et al., 2021). Thus, LRP1-mediated uptake of extracellular Tau may be a physiologic process; however, Tau in extracts from Alzheimer’s Disease brains partially escapes lysosomal degradation, seeding new intracellular aggregates (Rauch et al., 2020; Cooper et al., 2021).

Specific LRP1 ligands, including tissue-type plasminogen activator (tPA), α_2_M, and S-PrP^C^, activate anti-inflammatory cell-signaling responses in macrophages by a pathway that requires the NMDA-R (Mantuano et al., 2017, 2022). Determining whether Tau initiates LRP1-activated cell-signaling is an important question. There is considerable evidence that Tau accumulation in the CNS is pro-inflammatory (Yoshiyama et al., 2007; Nilson et al., 2017). In P301S-Tau transgenic mice, which express a mutated form of human Tau that forms intracellular filamentous lesions, microglial activation and neuro-inflammation precede formation of detectable Tau aggregates (Yoshiyama et al., 2007). Anti-inflammatory treatment blocks disease progression in P301S-Tau transgenic mice. Although activated microglia demonstrate some activities that are beneficial in AD, excessive microglial activation and chronic neuro-inflammation may promote AD progression (Akiyama et al., 2000; Heneka et al., 2015; Hong et al., 2016; Ransohoff, 2016).

In this investigation, we studied the 2N4R variant of Tau. Our findings indicate that Tau elicits a pro-inflammatory response in macrophages, microglia, and astrocytes in a manner entirely dependent on LRP1. Unlike other LRP1 ligands that activate NMDA-R-dependent pathways, the pro-inflammatory effects of Tau were NMDA-R-independent. These insights provide a new understanding of Tau’s role in AD pathology and open the door to therapeutic strategies that target Tau-mediated inflammation independently of NMDA-R.

## MATERIALS AND METHODS

### Proteins and reagents

Recombinant Human 2N4R Tau, the largest Tau derivative with 441 amino acids, was from R&D Systems. The NMDA-R inhibitors, dizocilpine (MK-801) and dextromethorphan hydrobromide (DXM), were from Cayman Chemicals and Abcam, respectively.

### Experimental mice

Wild-type (WT) C57BL/6J mice were obtained from Jackson Laboratory. Mice in which macrophages are LRP1-deficient were obtained by breeding *Lrp1*^flox/flox^ mice with mice express Cre recombinase under the control of the lysozyme-M (LysM-Cre) promoter, in the C57BL/6J background, as previously described (Overton et al., 2007; Staudt et al., 2013; Mantuano et al., 2016). To generate mice in which macrophages are deficient in the essential NMDA-R GluN1 subunit, *Grin1*^flox/flox^ mice were bred with mice that express Cre recombinase under the control of the LysM-Cre promoter in the C57BL/6J background (Mantuano et al., 2023). All research protocols involving mice were approved by the University of California San Diego Institutional Animal Care and Use Committee.

### Cell culture model systems

Bone-marrow-derived macrophages (BMDMs) were harvested from 16-week-old mice, as previously described (Mantuano et al., 2016). Briefly, bone marrow cells were flushed from mouse femurs, plated in non-tissue culture-treated dishes, and cultured in DMEM/F-12 medium containing 10% FBS and 20 ng/mL mouse macrophage colony-stimulating factor (BioLegend) for 7 days. Non-adherent cells were removed by washing. Adherent cells included >95% BMDMs, as determined by F4/80 and CD11b immunoreactivity.

Microglia were isolated from C57BL/6J mouse pups, as previously described (Ni and Aschner, 2010; Pontecorvi et al., 2019). In brief, brains were harvested from postnatal day 1–6 mice. The cortices were dissected from the forebrain, and the surrounding meninges were removed. Intact cortices were mechanically and enzymatically dissociated using the Neural Tissue Dissociation Kit (Miltenyi Biotec). Mixed glial cell cultures were established in DMEM/F-12 supplemented with GlutaMAX, 10% FBS, and 1× Gibco Antibiotic-Antimycotic (Thermo Fisher Scientific). After culturing for 10-14 days, microglia were dissociated by shaking the mixed cultures at 200 rpm for 30 min at 37°C. The floating cells were collected by centrifugation (5 min, 600×*g*) and re-plated at 3×10^5^ cells/well. Oligodendrocytes were removed by an additional 6 h of shaking. The remaining cells, which included mainly astrocytes, were collected by trypsinization and re-plated at 4×10^5^ cells/well. Experiments were performed within 24 h of completing the isolation procedure for microglia and within 48 h of completing the isolation procedure for astrocytes. Microglial cell culture purity was >95% homogeneous as determined by immuno-fluorescence (IF) microscopy for Iba1 (positive), glial fibrillary acidic protein (GFAP) (negative), and OLIG1 (negative). Astrocyte cultures were >95% GFAP-immunopositive.

### mRNA expression studies

BMDMs, microglia, and astrocytes were transferred to serum-free medium (SFM) for 1 h. Some cultures were pre-treated with MK-801 (1 µM) or DXM (10 µM) for the final 30 min of the 1 h pre-incubation. The cells then were treated for 6 h with Tau, at the indicated concentrations, or vehicle (20 mM sodium phosphate, 150 mM NaCl, pH 7.4, PBS) in the presence of inhibitors, as indicated. RNA was isolated using the NucleoSpin RNA kit (Macherey-Nagel) and reverse-transcribed using the iScript cDNA synthesis kit (Bio-Rad). qPCR was performed using TaqMan gene expression products (Thermo Fisher Scientific). Primer-probe sets were as follows: *Gapdh* (Mm99999915_g1); *Tnf* (Mm00443258_m1); *Il6* (Mm00446190_m1); *Ccl3* (Mm00441259_g1); and *Cxcl2* (Mm00436450_m1). The relative change in mRNA expression was calculated using the 2^ΔΔCt^ method with *Gapdh* mRNA as a normalizer.

### Cell signaling/immunoblotting experiments

BMDMs, microglia, and astrocytes were transferred to SFM for 1 h and then treated for 1 h with Tau or vehicle as noted. Some cultures were pre-treated with MK-801 (1 µM) or DXM (10 µM) for the final 30 min of the 1 h pre-incubation, as noted.

Extracts of BMDMs, microglia, and astrocytes, were prepared in RIPA buffer (Sigma-Aldrich) supplemented with protease inhibitors and phosphatase inhibitors (Thermo Fisher Scientific). Equal amounts of cellular protein were separated by SDS-PAGE and transferred to PVDF membranes. The membranes were blocked with 5% nonfat dried milk and then incubated with primary antibodies from Cell Signaling Technology that recognize: phospho-ERK1/2 (catalog #9102); total ERK1/2 (catalog #4370); phospho-IκBα (catalog #2859); total IκBα (catalog #9242); and β-actin (catalog #3700). The membranes were washed and incubated with horseradish peroxidase-conjugated secondary antibody from Jackson ImmunoResearch (anti-rabbit: catalog #111-035-003; anti-mouse: catalog #115-035-003). Immunoblots were developed using Thermo Scientific SuperSignal West Pico PLUS Chemiluminescent Substrate (Thermo Fisher Scientific) and imaged using the Azure Biosystems c300 digital system. When immunoblots were re-probed, phospho-IκBα was detected first, followed by total IκBα, then p-ERK, total ERK1/2, and finally, β-actin. Presented results are representative of at least three independent experiments.

### Proteome profiler mouse cytokine arrays

BMDMs and microglia were transferred to SFM for 30 min and then treated with Tau or vehicle for 6 h, as indicated. Conditioned medium (CM) was collected and clarified by centrifugation. An equivalent amount of CM (1.0 ml for each condition) was incubated with each nitrocellulose membrane provided in the Proteome Profiler Mouse XL Cytokine Array Kit (R&D Systems). Membranes were developed following the instructions of the manufacturer.

### Statistical analysis

Statistical analysis was performed using GraphPad Prism 9.5.1. All results are expressed as the mean ± SEM. Each replicate was performed using a different BMDM, microglia, or astrocyte preparation. Comparisons between two groups were performed using two-tailed paired t-tests. To compare more than two groups, we performed a one-way ANOVA followed by post-hoc Dunnett’s multiple comparison test. When evaluating the effects of two independent factors, we performed a two-way ANOVA followed by Šidák’s multiple comparisons test. *p*-values of \**p*<0.05, \*\**p*<0.01, \*\*\**p*<0.001, and \*\*\*\**p*<0.0001 were considered statistically significant.

## RESULTS

### Tau elicits an LRP1-dependent pro-inflammatory response in macrophages

BMDMs were treated with increasing concentrations of Tau for 6 h. RNA was harvested and RT-qPCR was performed to examine mRNA levels for the genes encoding various cytokines. Tau, at concentrations greater than or equal to 5 nM, significantly increased expression of the mRNAs encoding TNFα (*Tnf*), interleukin-6/IL-6 (*Il6*), CCL3/MIP-1α (*Ccl3*), and CXCL2/ MIP-2α (*Cxcl2*) (Fig. 1*A*).

**Figure 1.**
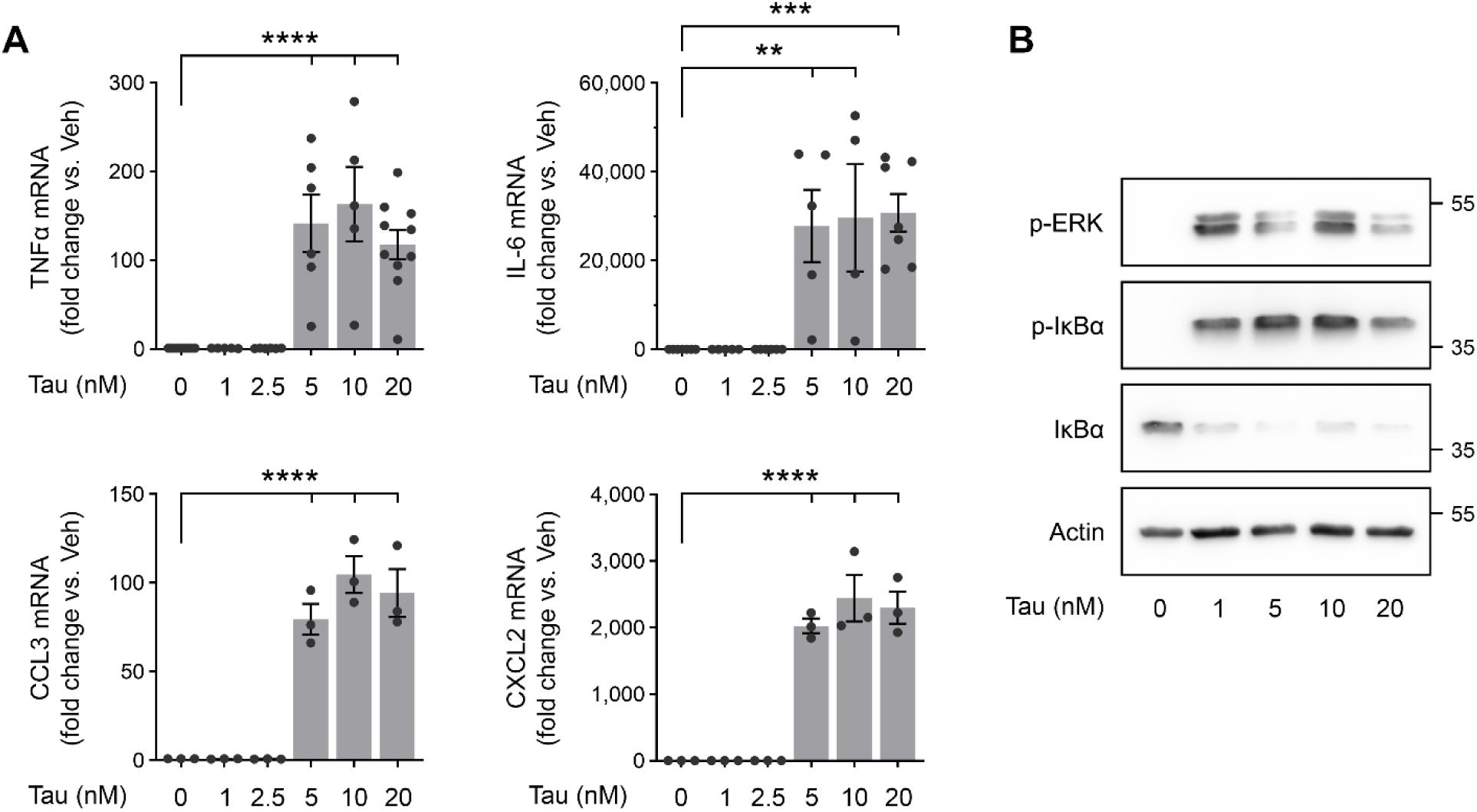
Tau induces an inflammatory response in macrophages. ***A***, BMDMs from wild-type mice were treated for 6 h with increasing concentrations of Tau, as indicated. RT-qPCR was performed to determine mRNA levels for TNFα, IL-6, CCL3, and CXCL2. Data are represented as mean ± SEM (n = 3-8; one-way ANOVA: \*\**p*<0.01; \*\*\**p*<0.001; \*\*\*\**p*<0.0001). ***B***, BMDMs from wild-type mice were treated for 1 h with increasing concentrations of Tau, as indicated. Immunoblot analysis was performed to detect expression of p-ERK, p-IκBα, IκBα, and β-actin.

IκBα phosphorylation is a key step in the activation of NFκB. Phosphorylated IκBα is ubiquitinated and degraded, allowing NFκB to translocate to the nucleus and bind DNA (Karin and Ben-Neriah, 2000; Viatour et al., 2005). NFκB activation promotes expression of pro-inflammatory cytokines, such as TNFα and IL-6. When BMDMs were treated for 1 h with Tau, at concentrations of 5 nM or greater, IκBα was phosphorylated and a decrease in the total abundance of IκBα was observed (Fig 1*B*). ERK1/2 also was phosphorylated in Tau-treated BMDMs. ERK1/2 phosphorylation is a commonly observed process in LRP1-initiated cell-signaling responses (Gonias and Campana, 2014).

To test whether LRP1 is necessary for Tau-induced cytokine expression, we harvested BMDMs from mice in which *Lrp1* is deleted under the control of the LysM promoter (Overton et al., 2007; Staudt et al., 2013; Mantuano et al., 2016). LRP1 protein is decreased by greater than 95% in these cells (Staudt et al., 2013). In LRP1-expressing control cells, which carry two floxed *Lrp1* genes but not LysM-Cre, 20 nM Tau significantly increased expression of the mRNAs encoding TNFα, IL-6, CCL3, and CXCL2, as anticipated (Fig. 2A). In LRP1-deficient BMDMs, 20 nM Tau failed to induce expression of these cytokines.

**Figure 2.**
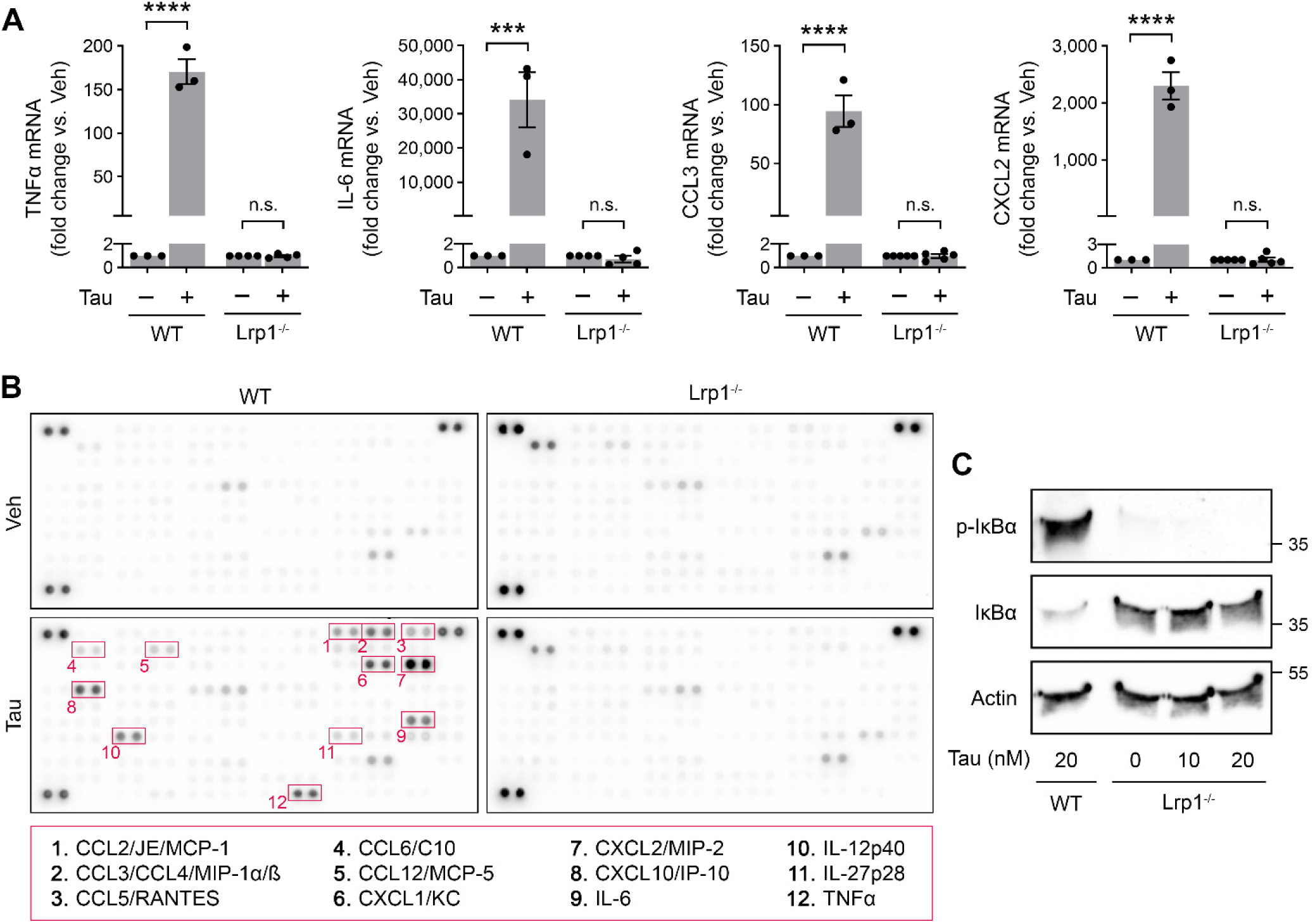
LRP1 mediates Tau-induced inflammation in macrophages. ***A***, BMDMs from wild-type (WT) and *Lrp1*^fl/fl^-LysM-Cre-positive (*Lrp1*^−/-^) mice were treated for 6 h with Tau (20 nM) or vehicle. mRNA levels for TNFα, IL-6, CCL3, and CXCL2 were determined by RT-qPCR. Data are represented as mean ± SEM (n = 3-5; two-way ANOVA: \*\*\**p* < 0.001; \*\*\*\**p* < 0.0001; n.s., not significant). ***B***, WT and *Lrp1*^−/-^ BMDMs were treated for 6 h with Tau (20 nM) or vehicle. Conditioned medium (CM) was collected and analyzed using the Proteome Profiler Mouse XL Cytokine Array. Representative cytokines that were increased in Tau-treated WT BMDMs compared with *Lrp1*^−/-^ cells are highlighted in red boxes and numbered. ***C***, WT and *Lrp1*^−/-^ BMDMs were treated for 1 h with Tau, at the indicated concentrations. Protein extracts were analyzed by immunoblotting to detect p-IκBα, total IκBα, and β-actin.

To examine cytokine expression at the protein level, in an unbiased manner, we treated LRP1-expressing and -deficient BMDMs with 20 nM Tau for 6 h. Conditioned medium (CM) was harvested and subjected to array analysis to detect secreted cytokines. In CM from Tau-treated LRP1-expressing cells, we observed TNFα and IL-6, as anticipated based on our RT-qPCR results (Fig. 2*B*). Other cytokines that were increased in CM from Tau-treated LRP1-expressing cells included but were not limited to CXCL2/MIP2, CXCL1/KC, CXCL10/ IP10, CCL3/CCL4 (MIP1α/β), and IL-12. By contrast, CM from Tau-treated LRP1-deficient BMDMs was entirely lacking in the Tau-elicited cytokines detected by this array. In separate studies, Tau failed to induce phosphorylation of IκBα in LRP1-deficient BMDMs, even when the Tau concentration was increased to 20 nM (Fig. 2*C*). Collectively, these results demonstrate that LRP1 is necessary for Tau-induced pro-inflammatory responses in BMDMs.

### The NMDA-R is not required for macrophages to respond to Tau

tPA, α_2_M, and S-PrP^C^ are examples of LRP1 ligands that activate cell-signaling via pathways that require the NMDA-R (Bacskai et al., 2000; Martin et al., 2008; Mantuano et al., 2017, 2020). To test whether the NMDA-R is necessary for Tau-elicited responses in BMDMs, first we treated cells with two pharmacologic inhibitors of the NMDA-R. Neither MK-801 nor DXM inhibited the effects of Tau on expression of the mRNAs encoding TNFα or IL-6. (Fig. 3*A*). Similarly, Tau-induced IκBα phosphorylation was unchanged in the presence of MK-801 or DXM (Fig. 3*B*).

**Figure 3.**
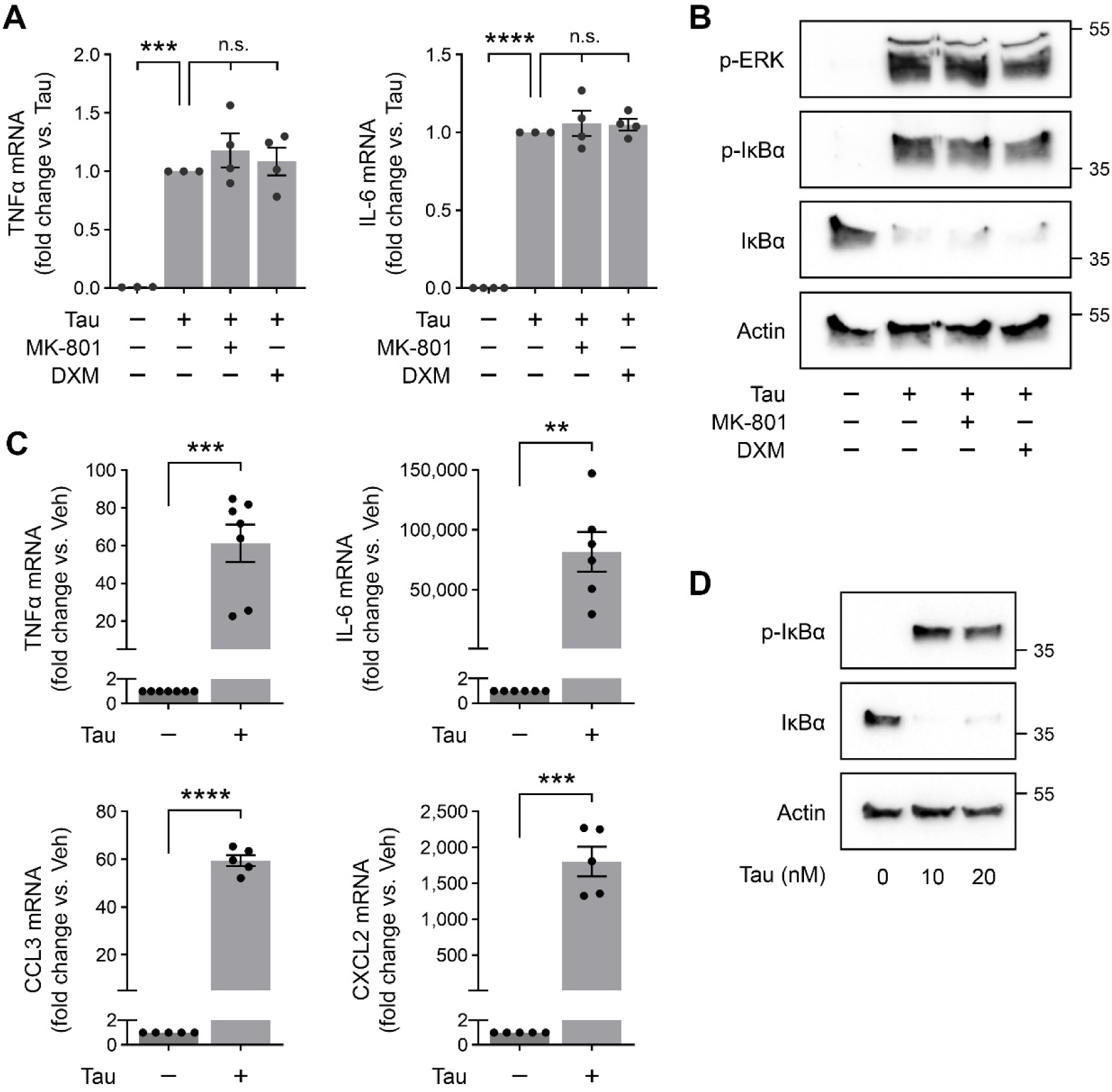
The Tau response in macrophages is independent of the NMDA-R. ***A***, BMDMs from wild-type (WT) mice were treated for 6 h with Tau (20 nM), MK-801 (1 µM), DXM (10 µM), or vehicle as indicated. RT-qPCR was performed to determine mRNA levels for TNFα and IL-6 (mean ± SEM; n=3-4; one-way ANOVA: \*\*\**p*<0.001; \*\*\*\**p*<0.0001; n.s., not significant). ***B***, WT BMDMs were treated for 1 h with Tau (20 nM), MK-801 (1 µM), DXM (10 µM), or vehicle as indicated. Cell extracts were subjected to immunoblot analysis to detect p-ERK, p-IκBα, total IκBα, and β-actin. ***C***, BMDMs from *Grin1*^fl/fl^-LysM-Cre-positive (*Grin1*^−/-^) mice were treated for 6 h with Tau (20 nM) or vehicle. mRNA levels for TNFα, IL-6, CCL3, and CXCL2 were determined by RT-qPCR (mean ± SEM; n = 5-7; paired t-test: \*\**p*<0.01; \*\*\**p*<0.001; \*\*\*\**p*<0.0001). ***D***, *Grin1*^−/-^ BMDMs were treated for 1 h with increasing concentrations of Tau, as indicated. Immunoblot analysis was performed to detect p-IκBα, IκBα, and β-actin.

Next, we isolated NMDA-R-deficient BMDMs from mice in which the gene encoding the essential GluN1 subunit (*Grin1*) is deleted under the control of the LysM promoter (Mantuano et al., 2023). BMDMs from these mice fail to respond to S-PrP^C^ (Mantuano et al., 2023). However, as shown in Fig. 3*C*, 20 nM Tau significantly increased expression of the mRNAs encoding TNFα, IL-6, CCL3, and CXCL2 in NMDA-R-deficient BMDMs. Furthermore, Tau (10 nM and 20 nM) caused IκBα phosphorylation in NMDA-R-deficient BMDMs, together with the concomitant decrease in total abundance of IκBα (Fig. 3*D*). Taken together, these results demonstrate that Tau-activated cell-signaling in BMDMs occurs independently of the NMDA-R.

### Tau induces LRP1-dependent pro-inflammatory responses in microglia and astrocytes

Microglia isolated from mouse pups were treated with 20 nM Tau or vehicle for 6 h. CM was harvested and subjected to cytokine array analysis (Fig. 4*A*). Multiple cytokines were detected, in a pattern that overlapped with those detected in Tau-treated BMDMs, including TNFα, IL-6, CCL3/CCL4, and CXCL2. To validate the results of the microarray analysis and test the role of the NMDA-R in Tau-elicited cytokine expression in microglia, we performed RT-qPCR studies. Tau (20 nM) increased expression of the mRNAs encoding TNFα, IL-6, CCL3, and CXCL2 in microglia (Fig. 4*B*). The increases in mRNA expression were not altered by the NMDA-R inhibitors, MK-801 and DXM.

**Figure 4.**
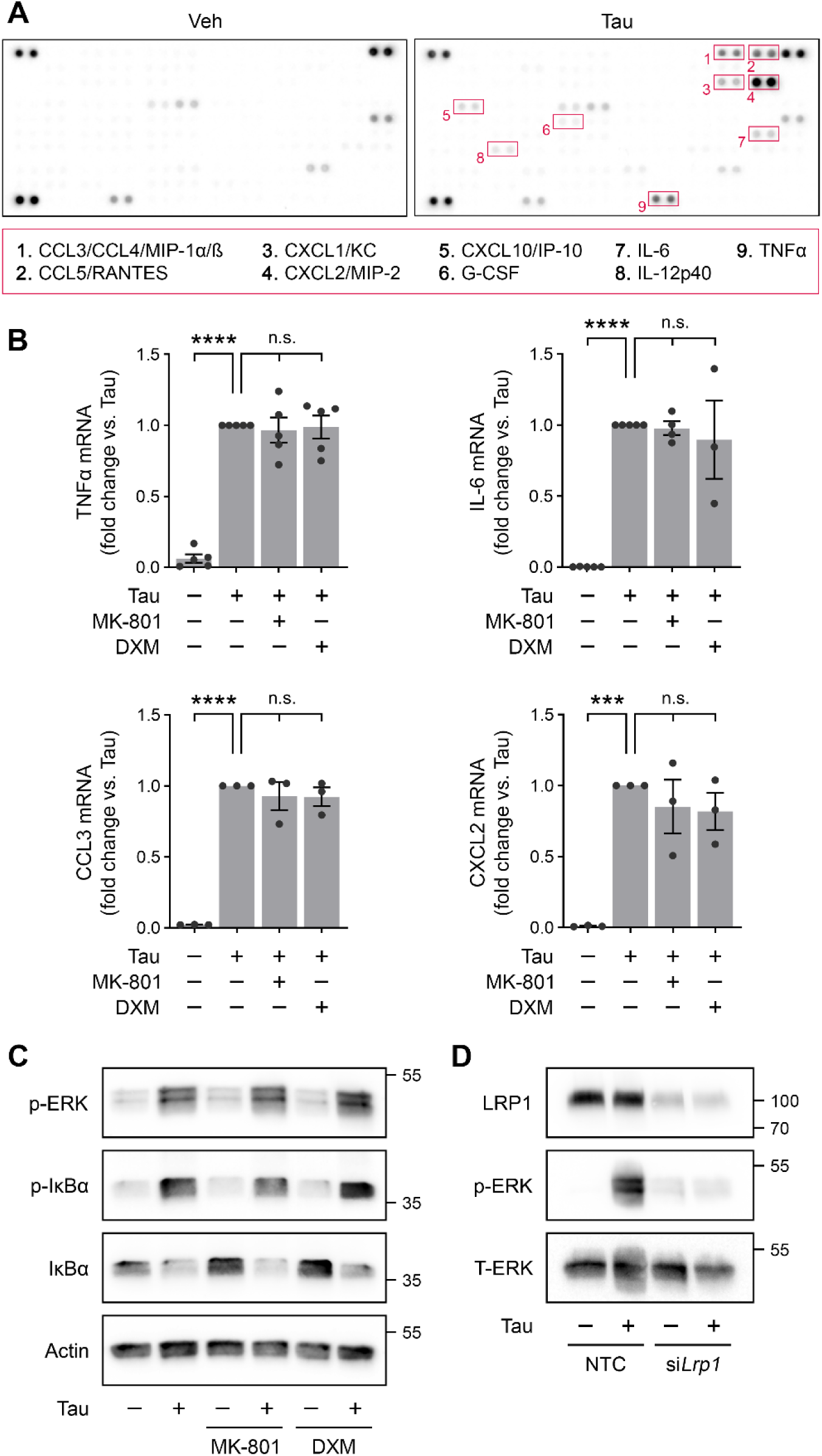
Tau induces a pro-inflammatory response in microglia that requires LRP1, but not the NMDA-R. ***A***, Microglia from mouse pups were treated with Tau (20 nM) or vehicle for 6 h. Conditioned medium (CM) was collected and analyzed using the Proteome Profiler Mouse XL Cytokine Array. Representative cytokines that were increased compared with the control are highlighted in red boxes and numbered. ***B***, Microglia from mouse pups were treated for 6 h with Tau (20 nM), MK-801 (1 µM), DXM (10 µM), or vehicle as indicated. RT-qPCR was performed to determine mRNA levels for TNFα, IL-6, CCL3, and CXCL2 (mean ± SEM; n = 3-5; one-way ANOVA: \*\*\**p*<0.001; \*\*\*\**p*<0.0001; n.s., not significant). ***C***, Microglia from mouse pups were treated for 1 h with Tau (20 nM), MK-801 (1 µM), DXM (10 µM), or vehicle as indicated. Cell extracts were subjected to immunoblot analysis to detect p-ERK, p-IκBα, IκBα, and β-actin. ***D,*** Microglia from mouse pups were transfected with *Lrp1*-specific (siL*rp1*) or non-targeting control (NTC) siRNA for 48 h. Cells were then treated with Tau (20 nM) or vehicle for 1 h and then analyzed by immunoblotting to detect LRP1, p-ERK, and T-ERK.

ERK1/2 was activated in microglia treated with 20 nM Tau for 1 h (Fig. 4*C*). In the same cells, IκBα was phosphorylated and this event was accompanied by a decrease in the total abundance of IκBα. These cell-signaling events were not inhibited by MK-801 and DXM, confirming that the NMDA-R does is not required for the response to Tau in microglia.

To test the role of LRP1 as a mediator of the Tau response in microglia, we silenced expression of *Lrp1* using siRNA. The level of LRP1 was substantially decreased at the protein level, compared with cells transfected with NTC siRNA (Fig. 4*D*). Tau (20 nM) failed to activate ERK1/2 in microglia in which *Lrp1* was silenced.

In cultured mouse astrocytes, Tau (20 nM) induced expression of the mRNAs encoding TNFα, IL-6, CCL3, and CXCL2 (Fig. 5*A*). The response to Tau in astrocytes was not inhibited by MK-801 or DXM. Tau also caused IκBα phosphorylation in astrocytes (Fig. 5*B*). This response was not inhibited by MK-801. Collectively, these results demonstrate that extracellular 2N4R Tau elicits similar pro-inflammatory responses in macrophages, microglia and astrocytes by an LRP1-dependent mechanism.

**Figure 5.**
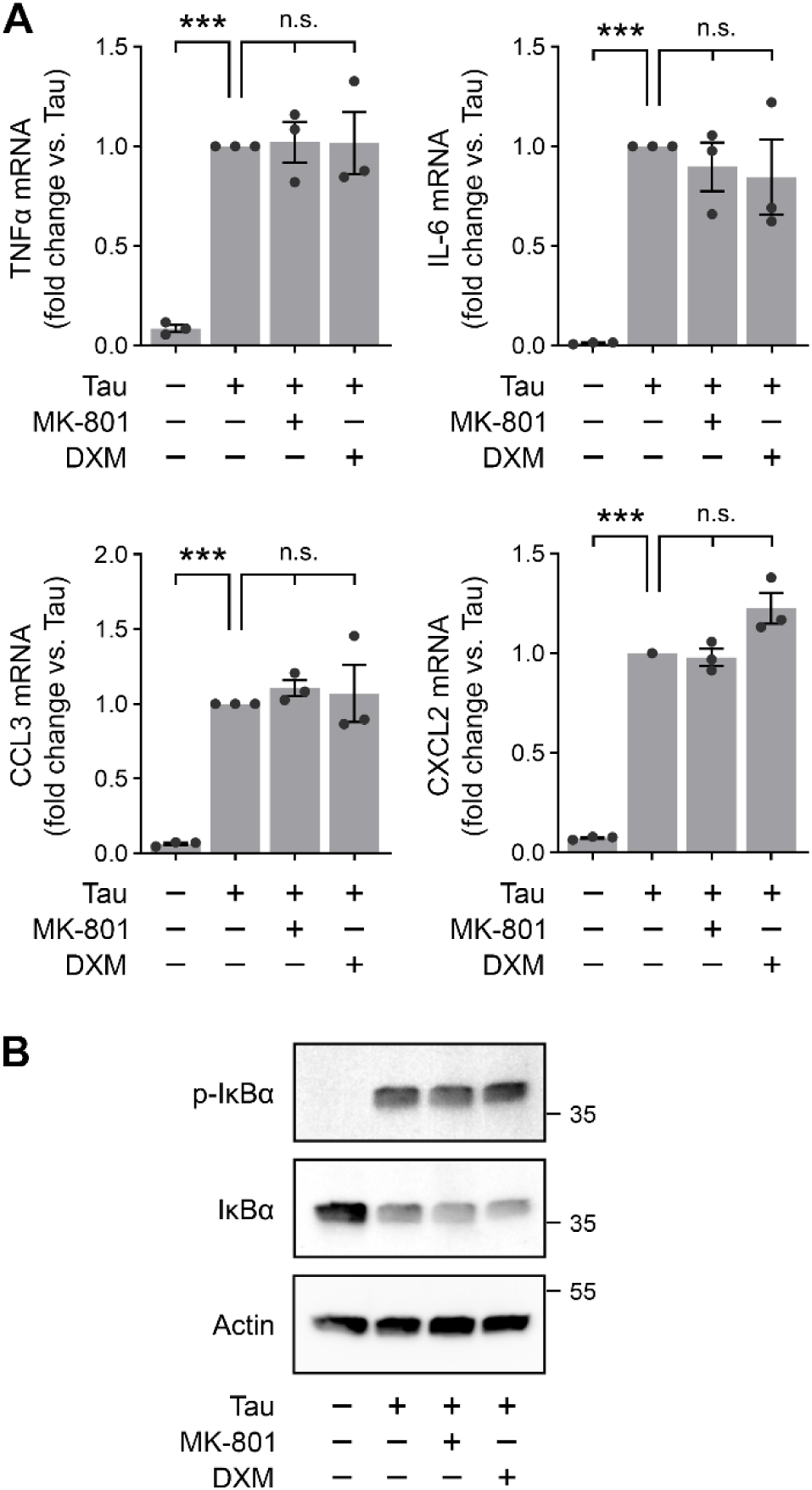
Tau induces a pro-inflammatory response in astrocytes that does not require the NMDA-R. ***A***, Astrocytes from mouse pups were treated for 6 h with Tau (20 nM), MK-801 (1 µM), DXM (10 µM), or vehicle as indicated. RT-qPCR was performed to determine mRNA levels for TNFα, IL-6, CCL3, and CXCL2 (mean ± SEM; n = 3; one-way ANOVA: \*\*\**p* < 0.001; n.s., not significant). ***B***, Astrocytes from mouse pups were treated for 1 h with Tau (20 nM), MK-801 (1 µM), DXM (10 µM), or vehicle as indicated. Protein extracts were analyzed by immunoblot to detect levels of p-IκBα, IκBα, and β-actin.

## DISCUSSION

LRP1 functions in the clearance of proteins from extracellular spaces, in lipoprotein uptake, as a regulator of the plasma membrane proteome, in myelin phagocytosis, in efferocytosis, and as a direct activator of cell-signaling (Strickland et al., 2002; Gonias and Campana, 2014). In AD, LRP1 may regulate disease progression by functioning as a receptor for β-amyloid peptide (Aβ) (Kanekiyo et al., 2013). LRP1 binds secreted forms of amyloid precursor protein (APP) and may regulate processing of cellular APP so that the amount of Aβ generated and secreted by cells is altered (Kounnas et al., 1995; Ulery et al., 2000). Isoforms of apolipoprotein E bind to LRP1 (Liu et al., 2007). Furthermore, LRP1 has been implicated in transport of Aβ across the blood-brain barrier (Deane et al., 2009).

The identification of LRP1 as a Tau receptor raises new questions regarding the activity of LRP1 in AD and in Tauopathies (Rauch et al., 2020; Cooper et al., 2021). Because binding of Tau to LRP1 does not require post-translational modification events such as phosphorylation or aggregation (Cooper et al., 2021), LRP1 may play an important physiologic role, removing low levels of Tau form the extracellular spaces in the CNS and delivering the internalized Tau to lysosomes for degradation. However, because Tau may partially escape endosomal transport and enter the cytoplasm, when the amount of Tau in the extracellular spaces in the CNS increases, LRP1 may be involved in propagation of Tau lesions in new cells (Rauch et al., 2020; Cooper et al., 2021).

Because LRP1 is known to couple ligand internalization with activation of cell-signaling (Gonias and Campana, 2014), our goal was to test whether the Tau-LRP1 interaction affects cell physiology in a manner that may play a role in progression of AD and other forms of neurodegeneration. We have shown that, in diverse cells, including macrophages, microglia, and astrocytes, the interaction of 2N4R Tau with LRP1 induces pro-inflammatory responses. Our study shows that these responses occur independently of the NMDA-R, which has been implicated in other LRP1 cell-signaling events (Bacskai et al., 2000; Martin et al., 2008; Mantuano et al., 2017, 2020).

Given that the NMDA-R has been implicated in the pathology of AD through its role in excitotoxicity and inflammatory signaling, the finding that Tau-LRP1 signaling is NMDA-R-independent raises questions about the co-receptors or alternative signaling pathways that might mediate Tau’s pro-inflammatory effects. Recent studies have suggested that other receptor systems, such as Toll-like receptor 4 (TLR4), may be involved in Tau-induced neuroinflammation, providing a potential explanation for how Tau can trigger inflammation independently of NMDA-R (Calvo-Rodriguez et al., 2020). Further studies are needed to explore whether TLR4 or other receptors act in concert with LRP1 to propagate Tau-driven inflammation in AD.

LRP1’s role in Tau clearance and pro-inflammatory signaling makes it a promising therapeutic target for AD. Modulating LRP1 activity could have dual benefits: promoting the clearance of extracellular Tau to reduce its propagation while inhibiting the pro-inflammatory signaling that exacerbates neurodegeneration. Because LRP1-mediated Tau signaling occurs independently of NMDA-R, targeting LRP1 may allow for more specific interventions that do not interfere with NMDA-R’s essential functions in synaptic transmission. This approach could avoid the limitations of current NMDA-R-targeting drugs like memantine, which aim to reduce excitotoxicity but may not address the full scope of Tau’s neurotoxic effects (Kabir et al., 2019). Another critical aspect of LRP1 biology is its ability to undergo proteolytic shedding. In response to inflammatory stimuli, such as LPS or interferon-γ, LRP1 is cleaved by transmembrane proteases in the ADAM family, releasing a soluble product that includes the entire α-chain and a fragment of the β-chain ectodomain (Quinn et al., 1999; von Arnim et al., 2005; Gorovoy et al., 2010). Shed LRP1 is found in plasma and its level is increased in patients with rheumatoid arthritis or systemic lupus erythematosus (Gorovoy et al., 2010). Shed LRP1 that is purified from human plasma is biologically active and independently pro-inflammatory against macrophages and microglia (Gorovoy et al., 2010; Brifault et al., 2017). It is possible that LRP1 shedding is involved in the pathway by which Tau induces pro-inflammatory responses in macrophages, microglia, and astrocytes. If future studies confirm this, it would suggest that shed LRP1 acts as a soluble factor that amplifies neuroinflammation by interacting with receptors on nearby cells. This could further potentiate the effects of Tau, promoting widespread inflammation and neurodegeneration in the CNS.

There are two possible models that could explain how LRP1 mediates Tau-induced inflammation. In the first model, membrane-anchored LRP1 functions with co-receptors to initiate Tau signaling at the cell surface. In the second model, shed LRP1 acts as a soluble factor, triggering pro-inflammatory responses by interacting with receptors on neighboring cells. These two models are not mutually exclusive, and both may contribute to the complex dynamics of Tau-induced neuroinflammation.

The bimodal activity of LRP1 as a ligand-dependent inhibitor or activator of inflammation is important to consider when reviewing complex animal model studies in which LRP1 may be exposed to diverse ligands. For example, LRP1 deficiency in cultured microglia is mildly pro-inflammatory in the absence of inflammation-regulating mediators (Yang et al., 2016; Brifault et al., 2017). LRP1-deficient microglia in culture demonstrate more robust pro-inflammatory responses to Aβ_1-42_ (He et al., 2020). In these cases, the effects of LRP1 deficiency may be explained by loss of NMDA-R-dependent anti-inflammatory cell-signaling pathways. By contrast, in an *in vivo* model of peripheral nerve injury, microglial LRP1 exacerbates inflammation in the spinal cord, in association with central sensitization of pain processing (Brifault et al., 2019). In this case, the available LRP1 ligands *in vivo* may activate the LRP1 pathway and/or induce LRP1 shedding. Animal models of AD should be carefully examined to determine how LRP1’s diverse functions influence disease progression, particularly in response to its various ligands.

In summary, the studies presented here identify a novel LRP1-depedent mechanism by which Tau may regulate inflammation, neuro-inflammation, and AD progression. The NMDA-R-independent nature of Tau’s pro-inflammatory effects opens new possibilities for targeting LRP1-Tau interactions as a therapeutic strategy. The potential involvement of shed LRP1 in amplifying Tau-induced inflammation highlights additional layers of complexity in LRP1 biology that warrant further exploration. By developing therapeutic strategies that modulate LRP1 activity, we may be able to slow AD progression by reducing Tau-induced neuroinflammation and preventing the spread of Tau pathology within the brain.

## Acknowledgements

This work was funded by a supplement (06S1) to NIH grant R01 HL136395.

## Author contributions

P.A., E.M., and S.L.G. designed the research; P.A., E.M., B.P., and C.Z. performed the experiments; all authors analyzed the data; P.A. performed statistical analyses and generated the figures; S.L.G. wrote the first draft of the paper with assistance from P.A. and E.M.; all authors approved the final draft of the paper.

